# Intranasal administration of *Burkholderia cepacia* promotes progressive acute inflammatory changes in experimental BALB/c mice

**DOI:** 10.64898/2026.05.26.727932

**Authors:** Selvaraj Roshan, Thippeswamy Aishwarya, Ramappa Chidananda, Inupanurthi Suma Madhuri, Muthuvel Atchaya, Abdul R Anshad, Sourabh R Bhujbal, D. Siva Sundara Kumar, Parthiban Rudrapathy, Pitchaipillai S. Ganesh, Sivadoss Raju, Suvaiyarasan Suvaithenamudhan, Ramendra Pati Pandey, Sridhar Muthusami, Rajesh Nachiappa Ganesh, Latchoumycandane Calivarathan, Esaki M Shankar

**Author notes:** **Corresponding authors:** Esaki M. Shankar, Department of Biotechnology, School of Integrative Biology, Central University of Tamil Nadu, Thiruvarur 610101, India. eMail, Latchoumycandane Calivarathan, Department of Biotechnology, School of Integrative Biology, Central University of Tamil Nadu, Thiruvarur 610101, India. These authors contributed equally (Selvaraj Roshan, Thippeswamy Aishwarya, Ramappa Chidananda).

## Abstract

*Burkholderia cepacia* (*B. cepacia*) is an opportunistic pathogen with versatile virulence mechanisms. The pathogenesis of *B*.*cepacia* in the immunocompetent host following intranasal exposure largely remains ambiguous. Male BALB/c mice were intranasally inoculated with *B. cepacia* strain 20209 (1×10□ CFU) and evaluated on days 3, 7, 14, and 21 post-infection. Histopathology of lung, liver, spleen, and kidney tissues were performed using H&E and PAS staining. Plasma cytokines were quantified using commercial multiplex assays and ELISA. Matrix metalloproteinase-2 (MMP-2) activity was assessed via gelatin zymography and metabolomic profiling by high-resolution mass spectrometry (HRMS). Histopathological analysis revealed organ-specific pathological indices such as interstitial pneumonitis, bronchitis, leukocyte infiltration, hepatic inflammation, as well as splenic hyperplasia. Similarly, MMP-2 activity revealed time-dependent modulation, reflecting dynamic proteolytic responses. Plasma and tissue IL-18 and IL-1β levels demonstrated a temporal regulation, with IL-18 peaking on day 7 post-infection, while IL-1β showed a biphasic expression peaking on day 3 and 14. Untargeted metabolomics revealed differential expression of lipid metabolism, and energy pathways, with higher expression of phospholipids and sphingolipids. Together, our study portrayed a physiologically relevant intranasal BALB/c model that captures both localized and systemic inflammatory responses to *B. cepacia*. Our findings highlight organ-specific pathologic progression and sustained inflammation providing key insights into host–pathogen interactions.

## INTRODUCTION

*Burkholderia cepacia* (*B. cepacia*) is a Gram-negative, motile, aerobic, non-fermenting, oxidase- and catalase-positive bacterium implicated with opportunistic infections, especially in individuals with compromised immunity (Avgeri et al., 2009). *B. cepacia* is ubiquitously distributed in nature, especially in plants, as a phytopathogen (Jin *et al*., 2020). *B. cepacia* complex (Bcc) includes several genetically distinct species and genomovars (Speert, 2002). Furthermore, nine distinct groups within the Bcc have been described based on DNA sequencing and genome-based analysis, demonstrating their complex genetic diversity and taxonomic heterogeneity (Sfeir, 2018). Members of the Bcc primarily afflict individuals with underlying cystic fibrosis (CF) or chronic granulomatous disease (CGD) (Gautam et al., 2011). In CF patients, CFTR mutations impair mucociliary clearance, promoting chronic lung colonisation by *B. cepacia* (Parameswaran and Murphy, 2007). Moreover, *B. cepacia* infection leads to a sharp decline in lung functions, increased morbidity, and risk of onset of “*cepacia syndrome*,” marked by bacteremia (Karakoc Parlayan et al., 2025; Tavares et al., 2020). CGD patients become increasingly vulnerable to invasive *B. cepacia* infection owing to defective oxidative burst, as the bacterium can effectively circumvent non-oxidative intracellular killing (Karakoc Parlayan *et al*., 2025). Several clinical studies involving CF and CGD confirm that adverse clinical outcomes are often associated with relatively reduced eligibility for lung transplantation and increased mortality (Chiarini et al., 2006).

*B*.*cepacia* displays numerous virulence and adaptive traits, including biofilm formation, quorum-sensing–mediated persistence, cell interaction, lipopolysaccharides (LPS) that augment resistance to host defences, and intracellular survival (Diban et al., 2023). *B. cepacia* shows high genomic flexibility and regulatory capacity, enabling rapid adaptation to conditions like low oxygen and immune pressure in the CF lung mucus (Sousa et al., 2017) (Hassan et al., 2019). Although *B. cepacia* is a known opportunistic pathogen in immunocompromised individuals, it’s pathogenesis in the healthy host remain elusive (Sousa et al., 2007). Moreover, the systemic host responses following intranasal exposure (which often represents their natural route of entry) remain obscure. Studies conducted to date have been contingent on intraperitoneal or aerosol challenge models and/or have used immunosuppression as a means to establish *B. cepacia* infection. Such experimental models may seldom represent natural disease acquisition and progression (D. P. Speert et al., 1999a; Vanhoutte et al., 2017).

BALB/c mice are widely used in experimental respiratory infections owing to their well-characterised immune system, and predictable susceptibility to pulmonary pathogens. Existing intranasal models fail to accurately mimic the systemic spread of *B. cepacia* and the accompanying hematological and organ-specific histopathological changes observed in humans. This paucity for a comprehensive, integrated host response atlas limits a thorough understanding of the immunopathogenesis of *B. cepacia* infection (Vanhoutte *et al*., 2017). A reliable experimental animal model mimicking the physiological responses of infected patients is critical to improved understanding of *B*.*cepacia* immunopathogenesis (Wisplinghoff, 2017). The development of a suitable animal model will facilitate the transition of animal data to clinical data, which fortifies the link between experimental and clinical research (D. P. Speert et al., 1999a). Here, we sought to explore the spatiotemporal immunohistopathological responses following intranasal exposure of BALB/c mice to *B. cepacia* strain 20209, to gain a deeper understanding of the ensuing inflammatory pathological consequences in the host.

## METHODS

### Ethics approval

All animal experiments were performed in accordance with the guidelines of the Committee for Control and Supervision of Experiments on Animals (CPCSEA), Government of India. The study protocol, including animal handling, acclimatisation, infection protocols/procedures, sampling schedule and end-point criteria, were reviewed and approved by the Institutional Animal Ethics Committee (IAEC) of Annamalai University, Chidambaram (Approval No.: GMCHC-IAEC/PR/1412/4/25).

### Animals

Male BALB/c mice of age 6-8 weeks and weighing 18-22 g were procured from Biogen Laboratory Animal Facility, Bangalore (a CPCSEA-registered breeder in India). Animals were housed in individually ventilated, sterile, isolated cages under specific pathogen-free (SPF) conditions, with controlled temperature, humidity, 12-hour light and 12-hour dark cycle. Mice were provided *ad libitum* access to food and water. All animals were acclimatised to the animal facility conditions before initiation of the experiments, and remained under the same conditions throughout the experimental time-period.

### Bacteria

*B. cepacia* (Strain No. 20209) was maintained as frozen stocks at −80°C in Luria-Bertini (LB) broth (Cat. No.: SKU G1245; Miller formulation; HiMedia Laboratories, Mumbai, India) supplemented with 25% (v/v) glycerol. The phenotypic characterization of the *B. cepacia* strain was confirmed as described previously (Bhujbal et al., 2026) (**Supplementary Table 1**). For the experimental procedure, bacteria were sub-cultured from frozen stocks into Luria–Bertani broth and incubated for 18–24 h at 37°C (Gharat *et al*., 2025).

### Inoculum preparation

*B. cepacia* 20209 was cultured in LB broth (Miller) at 37°C for 16–18 h with shaking at 150 rpm, and maintained on LB agar (1.5%) (Cat. No.: SKU G1151; Miller formulation; HiMedia Laboratories, Mumbai, India). A single colony was picked from a freshly streaked LB plate and suspended in LB broth before washing in sterile phosphate-buffered saline (PBS) for inoculum preparation, followed by resuspension in PBS (David P. Speert *et al*., 1999). Bacterial concentration was estimated by measuring the optical density at 600 nm (OD_600_) and by generating a known correlation between OD and CFU (Mariappan *et al*., 2011). The inoculum was prepared and serial dilutions were done to confirm that viable bacteria were obtained. The suspension was maintained on ice until use in the experiments (U. Sajjan et al., 2001).

### Experimental groups and study design

The study design consisted of four experimental groups, each containing eight mice. In every group, six mice were intranasally inoculated with five microliters of *B. cepacia* 20209 inoculum (1 × 10^6^ CFU), and two mice served as controls and received only sterile PBS. Following inoculation, the animals were monitored daily. At pre-determined time-points (3, 7, 14, and 21 days post-infection), mice from the respective experimental groups were euthanized by cervical dislocation, and blood and organ samples were collected under aseptic conditions for experimental analyses.

### Sample collection

Post-infection, the mice in the respective groups were euthanised in accordance with established ethical guidelines on days 3, 7, 14, and 21. Blood was collected from euthanised mice under aseptic conditions in commercial BD Vacutainer K_2_EDTA blood collection tubes (Cat. No.: 368841; Becton Dickinson, Franklin Lakes, NJ, USA). The plasma was collected, and was aliquoted and stored at −80°C (Hassis *et al*., 2015). Organs such as liver, spleen, kidney and lungs were harvested and stored in 10% neutral buffered formalin (NBF) to preserve cellular architecture for histopathological examinations (Al-Sabaawy *et al*., 2021). Tissues were fixed and processed using standard procedures for paraffin embedding and staining (Shetty *et al*., 2020).

### Preparation of tissue lysate

To prepare liver tissue homogenates, frozen liver tissues stored at −80°C were disaggregated into small pieces and placed in pre-chilled microfuge tubes before adding a cocktail of 900 μL of PBS (Balkan et al., 2025), 100 μL protease inhibitor, and 20 μL phenylmethylsulfonyl fluoride (PMSF). The samples were homogenised using a tissue homogeniser until a uniform suspension was formed. After homogenization, the samples were vortexed for 20 seconds in five cycles, with each cycle followed by a two-minute interval between each cycle. Later, the samples were centrifuged at 14,000 x g for 15 mins at 4°C (Chien et al., 2025), and the supernatants were collected. The centrifugation process was repeated under the same conditions to yield a clear protein extract of tissue lysate. Samples of supernatants obtained from control and infected subjects on days 3, 7, 14, and 21 post-infection were stored for downstream analyses.

### Hematoxylin-Eosin (H&E) staining

Tissue sections, each measuring 4-5 μm in thickness, were prepared using a rotary microtome, and subsequently affixed to glass slides for microscopic examination. Following sectioning and mounting, all tissue samples were subjected to H&E staining to facilitate a histomorphological assessment of the overall structure and to observe if any pathological changes were linked to infection (Ah *et al*., 2008). Histopathological analysis was carried out using light microscopy to evaluate the inflammatory responses and to identify structural modifications in the primary organs. Representative samples were taken and the significant pathological findings, such as inflammatory cell infiltration, vascular congestion, necrosis, and the disorganization of normal tissue architecture were documented thereof. Photomicrographs of the respective tissues were captured using a digital imaging system to preserve the histopathological observations for further investigations (Rangan and Tesch, 2007).

### Periodic acid Schiff (PAS) staining

Tissue sections prepared were mounted on glass slides before treatment with 0.5 to 1% periodic acid solution for 5-10 min, rinsed in dH_2_O, followed by the addition of Schiff’s reagent for 10-15 min at room temperature. Slides were washed in running tap water for 5-10 min for the development of color. Counterstaining was performed using Mayer’s hematoxylin, followed by dehydration, clearing and mounting. Later, the slides were visualized using an RXLr-5NX series advanced research fluorescence microscope (Radical Scientific Equipment, Ambala, India).

### Multiplex cytokine assay

Cytokine IL-2, IL-4, IL-5, IL-10, GM-CSF, IFN-γ, and TNF-α levels in plasma were measured using a commercial Bio-Plex Pro Cytokine assay kit (Cat. No.: M600000007A; Bio-Rad, Hercules, CA, USA), according to the manufacturer’s instructions. Samples were diluted by 1:4 using sample diluent; analysed on the Bio-Plex 200 system (Cat. No.: 171000201; Bio-Rad Laboratories, Hercules, CA), and concentrations were calculated using Bio-Plex Manager Software (ver.6.0).

### Enzyme immunoassay

Enzyme-linked immunosorbent assay (ELISA) was performed using commercial high-sensitivity ELISA kits to quantify IL-1β (Cat. No.: E-HSEL-M0006-48T; Elabscience, Houston, TX, USA and IL-18 levels (Cat. No.: E-HSEL0M0001-48T; Elabscience, Houston, TX, USA) in plasma samples according to the manufacturer’s instructions. The samples were diluted in 1:2 ratio before measuring on an ELISA plate reader (BIO-RAD iMark microplate reader), and concentrations were calculated from the plotted standard curve.

### Gelatin zymography

Matrix metalloproteinase-2 (MMP-2) activity was assessed using gelatin zymography under non-reducing conditions. Prior to electrophoresis, plasma samples of control, day 3, 7, 14 and 21) were diluted at a ratio of 1:10 using sterile PBS. Samples were mixed with loading buffer without β-mercaptoethanol and were not heat-denatured (to preserve enzyme activity). Proteins were resolved on 10% SDS-PAGE gels co-polymerized with 0.1% gelatin as a substrate. Equal volumes of diluted serum samples were loaded into each well. Electrophoresis was performed at constant voltage until the dye front migration was complete. Following electrophoresis, gels were washed with wash buffer (Triton X-100, 50 mM Tris-HCl, pH 7.5, 5 mM CaCl_2_ and 1μM ZnCl_2_) for one hour to remove SDS. Subsequently, the gels were incubated in an activation and renaturation buffer containing (50 mM Tris-HCl, pH 7.5, 10 mM CaCl_2_ and 5μM Zncl_2_) at 37°C for 18–24 hours to facilitate enzymatic digestion of gelatin. After incubation, gels were stained with 0.5% Coomassie Brilliant Blue R-250 for 60 min and destained using a solution of methanol and acetic acid until clear bands appeared against a dark blue background. Proteolytic activity of MMP-2 was visualized as transparent bands corresponding to gelatin degradation zones.

### Coomassie brilliant blue staining

To verify equal protein loading, a parallel SDS-PAGE gel (without gelatin) was run under standard denaturing conditions. Serum samples (1:10 diluted) were mixed with reducing sample buffer, boiled at 95°C for five minutes, and loaded onto the gel. After electrophoresis, the gel was stained with Coomassie Brilliant Blue for 60 min, followed by destaining in methanol-acetic acid solution until clear protein band patterns were visible. The uniformity of total protein bands across lanes was used as internal loading control.

### High-resolution mass spectrometry

HR-MS is a highly sensitive and precise technique that enables comparative metabolite analysis between infected and control groups (Burlikowska *et al*., 2020). Serum samples were thawed on ice and processed for metabolite extraction. For HR-MS analysis, metabolites were isolated using an equal mixture of methanol and chloroform, following a slightly modified version of a previously reported protocol (McHugh et al., 2018). The mixture was vortexed, incubated at –20°C for 30 min, and then centrifuged at 12,000 rpm for 15 min. An aliquot of 20 μL of the supernatant was diluted with 980 μL of methanol for further metabolomic analysis. This method separates proteins besides isolating small metabolites present in the plasma (Gutmann *et al*., 2024). HR-MS analysis was performed using a Xevo G2-XS QTof mass spectrometer (Waters) in ES Unispray positive mode. The supernatant was diluted with methanol, and 1 μL was injected. Separation was carried out on an Acquity UPLC BEH C18 column (2.1 × 50 mm, 1.7 μm) at 120°C with a capillary voltage of 1 kV and collision energy of 10–30 V. Full-scan data were acquired over 100–1600 m/z, and metabolite identification was enabled by accurate mass measurements. Data were processed using MassLynx 4.2 software (Waters).

### Molecular docking

#### a) Preparation of ligand molecules

In order to prepare the selected compounds in an SDF file format, they were retrieved from the PubChem database. The compounds were then optimised using ligand preparation tools (to produce low-energy conformers) for subsequent use in the docking studies. Homology Modelling and Preparation of Protein Structure Hfq, porin-like protein, AHL receptor and acyl-homoserine-lactone synthase from *B. cepacia*, were homology modelled using the prime module available within the Schrödinger software suite. The three-dimensional (3D) structure for Hfq (PDB ID 3SB2_A, (Chain A, Protein hfq from *Herbaspirillum seropedicae* SmR1)), Porin 1 (PDB ID 5NXN_A, (Chain A, Porin 1 from *Providencia stuartii*)) and RhlR (PDB ID 7R3G_A, (Chain A, regulatory protein RhlR from *Pseudomonas aeruginosa* PAO1)). The other model ligand synthase was developed using trial 74% similarity amongst the template proteins with AHL synthase from *Burkholderia glumae* (PDB ID 3P2f_A, (Chain A, AHL synthase from *Burkholderia glumae*)). Additionally, the 3D structure of the RNA-binding protein in their FASTA accession sequence for each of the proteins (B4EAW4, Q8K9I8, Q9AM46 and Q9ZIU1) was retrieved from the UniProt database [40] and was used for modelling the 3D structure. The loops and structures were further refined with the ‘Protein Refinement module’ [24] of the Schrödinger software. The molecular structure was prepared with molecular modelling software, where hydrogen atoms were added, water was removed, and bond orders were corrected. The geometry of the protein was optimised to prepare for energy minimisation of the protein xP3. The energy minimisation before docking analysis was performed to stabilise the proteins.

#### b) Molecular docking analysis

The docking analysis was conducted using Schrödinger Glide. The ligands were placed into the active binding sites of the following proteins: RNA-binding protein (Hfq), porin-like protein (BUsg_347), AHL receptor, and AHL synthase. The following parameters were used to assess the affinity of the ligands for their respective proteins: docking scores; number of hydrogen bonds formed; length of hydrogen bonds; number of hydrophobic contacts; and number of electrostatic interactions.

### Statistical analysis

The mice’s body weight and analyte concentration were expressed as mean±SD. The statistical analysis of ELISA and BioPlex cytokine array was performed using a non-parametric Kruskal-Wallis test, followed by Dunn’s multiple comparison post hoc test. A p-value of <0.05 was considered statistically significant. The significant values were represented as *<0.05, **<0.01. All the statistical analyses were performed using PRISM software ver.6.0 (GraphPad, CA, USA).

## RESULTS

### No significant difference was evident in body weights among the experimental animals following exposure to *B. cepacia*

To ascertain whether *B. cepacia* exerted any measurable effect on the overall systemic health of the experimental animals, body weights of the BALB/c mice were recorded at regular intervals throughout the experimental duration. Body weight represents a non-invasive indicator of general health and disease burden in murine models. Comparative analysis revealed no significant differences in body weight between the infected and control groups at any of the time-points studied. These findings suggest that despite the onset of infection and the observed pathological changes, *B. cepacia* did not drive any systemic deterioration or weight loss in the experimental mice **(Supplementary Table 2)**.

### Progressive interstitial pneumonitis and bronchitis were noticed in the lungs of BALB/c mice following *B. cepacia* infection

Next, we set out to perform a comprehensive histopathological examination of the lung, liver, spleen, and kidney tissues to examine infection-induced tissue alterations at days 3, 7, 14, and 21 following *B. cepacia* infection. Tissue sections were stained using H&E to evaluate general morphology and inflammatory changes, and PAS staining to detect polysaccharides and mucopolysaccharide-rich structures. H&E-stained sections of lungs of *B. cepacia*-infected mice revealed moderate and progressively increasing inflammatory changes over time. The observed features were strongly suggestive of interstitial pneumonitis, including thickening of alveolar septa, infiltration of inflammatory cells within the interstitial spaces, and disruption of normal alveolar architecture (**Figure 1**). Furthermore, evidence of bronchitis was noticeable, as indicated by peribronchial infiltration of inflammatory cells together with epithelial alterations. The severity and magnitude of the pathological features appeared to increase from day 3 through 21 post-infection, suggesting a sustained and progressive pulmonary inflammatory response. In contrast, lung sections of control animals consistently exhibited normal histoarchitecture, with well-defined alveolar spaces, thin septa, and absence of inflammatory infiltrates or structural abnormalities.

**Figure 1.**
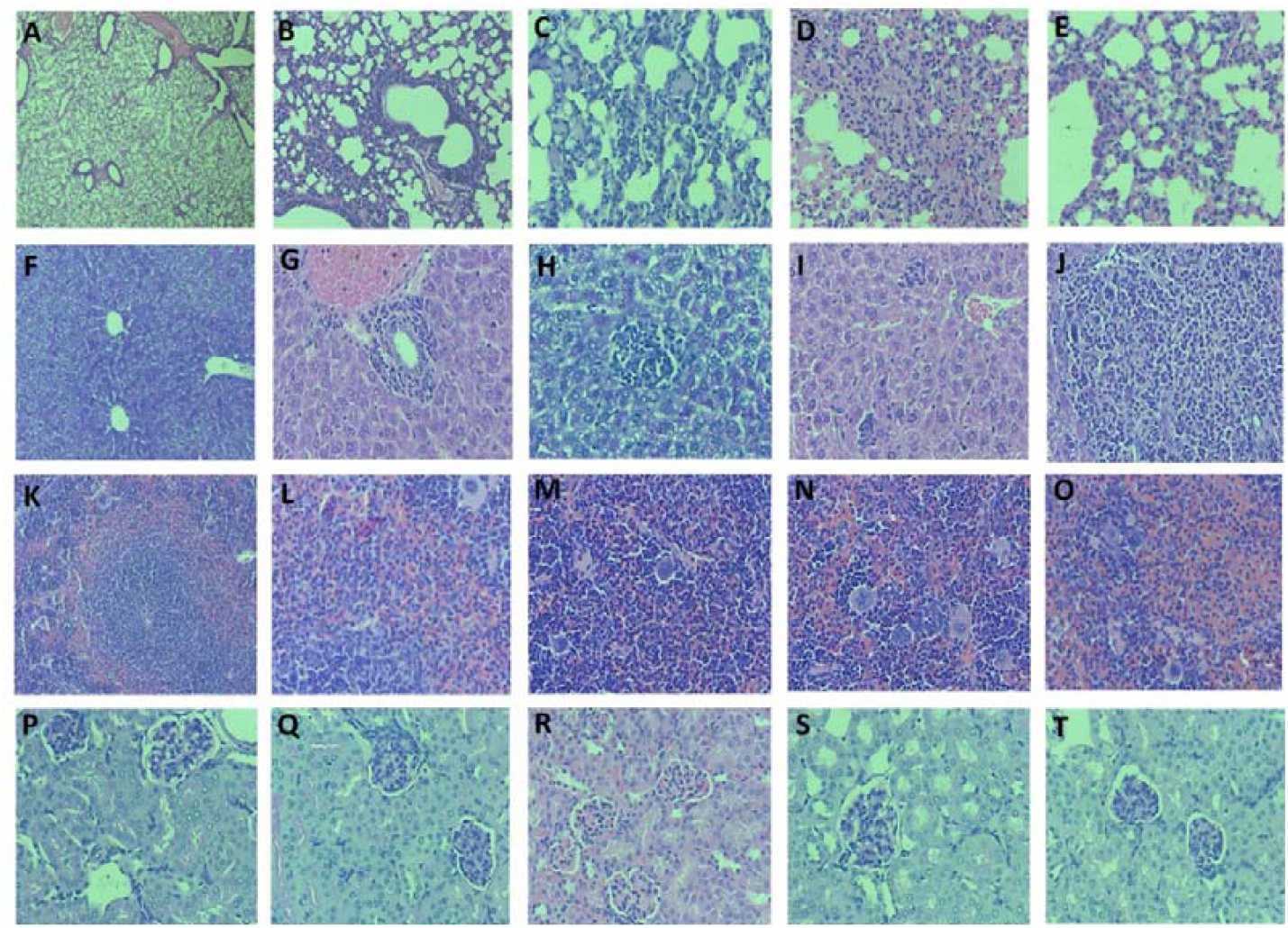
Histopathological staining of visceral organs of BALB/c mice intranasally administered with B.cepacia. **(A)** control lung section; **(B-E):** infected lung section (Day 3, 7, 14 & 21); **(F):** control liver section, **(G-J):** infected liver section (Day 3, 7, 14 & 21); **(K):** control spleen section; **(L-O):** infected spleen section (Day 3, 7, 14 & 21); **(P):** control kidney section; **(Q-T):** infected kidney section (Day 3, 7, 14 & 21). **(B-E)** shows increasing neutrophilic infiltration in lung parenchyma; **(G-J)** exhibits increasing spotty necrosis and microabscess in liver parenchyma; **(L-O)** shows expanding red pulp and dense neutrophilic infiltration in spleen section; **(Q-T)** shows no significant changes in the kidney section.

### *B. cepacia* infection resulted in progressive microabscess formation and focal liver necrosis in BALB/c mice

Next, we sought to investigate the histopathological aspects of liver sections of infected animals. We found progressively worsening pathological alterations in the liver tissue over time. The hallmark features included the formation of microabscesses, characterized by localized aggregates of inflammatory cells within the hepatic parenchyma, together with areas of spotty (focal) necrosis indicative of hepatocellular injury (**Figure 2**). These lesions were initially mild on day 3 post-infection but progressed to more pronounced lesions by days 7, 14, and 21, reflecting a gradual escalation in the severity of hepatic involvement. The progression appears to indicate sustained inflammatory activity and ongoing tissue damage in response to *B. cepacia* infection. In contrast, liver sections from control animals consistently exhibited normal histological architecture, with well-organized hepatic cords, intact hepatocytes, and clearly defined sinusoids (**Figure 1**). No evidence of inflammatory infiltration, necrosis, or structural disruption was observed at any of the examined time points, indicating the absence of pathological alterations in uninfected BALB/c mice.

**Figure 2.**
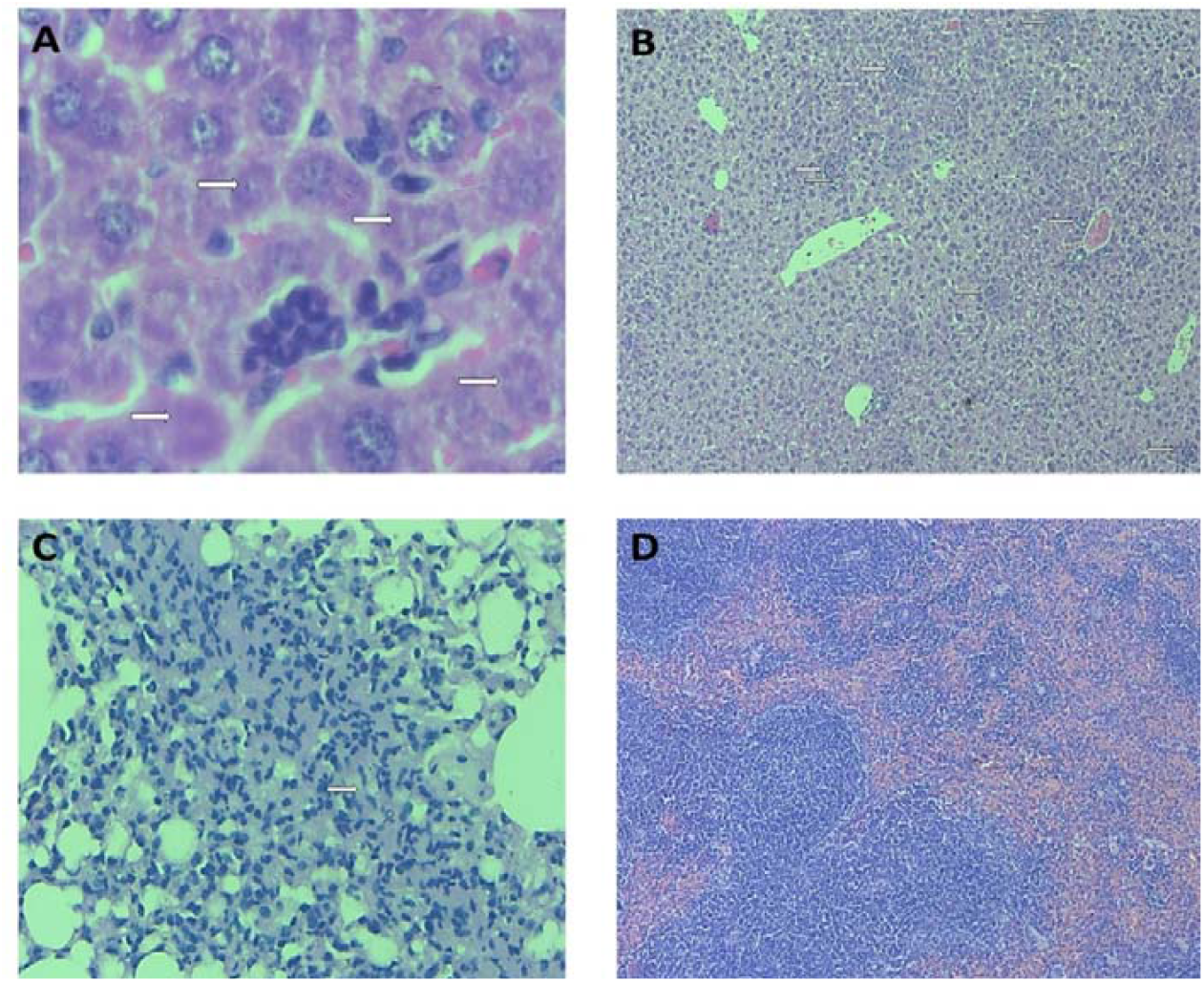
Histopathological staining of visceral tissues of BALB/c mice intranasally administered with B.cepacia. **(A)** Liver section - highlights foci of spotty necrosis of hepatocytes (arrows) evoking neutrophilic microabscesses (arrow head); **(B)** highlights liver parenchyma with multiple foci of spotty necrosis and microabscesses (arrows) randomly distributed in hepatic lobules; **(C)** highlights microgranuloma (arrow) surrounded by neutrophilic infiltration in both alveolar spaces and interstitium; **(D)** highlights spleen with massive expansion of red pulp.

### *B. cepacia* induced histopathological alterations in the spleen of BALB/c mice suggestive of red pulp expansion and progressive neutrophilic infiltration

Histopathological evaluation of spleen sections of infected animals demonstrated notable structural and cellular alterations that progressed across the infection course. A prominent finding was the expansion of the red pulp region, indicative of increased vascular congestion and heightened immune activity. Furthermore, there were heightened signs of neutrophilic infiltration, reflecting an active innate immune response to *B. cepacia*. The degree of red pulp expansion and density of infiltrating neutrophils increased progressively from day 3 through day 21 post-infection, suggesting sustained splenic involvement and an ongoing systemic inflammation. In contrast, spleen sections of control animals exhibited normal histoarchitecture, with well-defined white pulp and red pulp regions lacking any evidence of inflammatory cell infiltration or structural disruption (**Figure 2**). Together, our observations indicate that the changes seen in the *B. cepacia*-infected groups are attributable to infection-induced inflammation.

### *B. cepacia* exposure preserved the normal renal architecture in *B. cepacia*-infected mice and did not compromise renal integrity

Next, we examined the histopathological correlates of kidney sections of both *B. cepacia*-infected and control animals, which intriguingly revealed well-preserved renal architecture throughout the longitudinal study tenure. Across all time-points (days 3, 7, 14 & 21 post-infection), the kidneys displayed normal structural organization, including intact glomeruli, well-defined Bowman’s capsules, and properly arranged renal tubules with no evidence of cellular degeneration or necrosis. There were no signs of inflammatory cell infiltration, interstitial edema, vascular congestion, or tubular damage in the infected BALB/c mice. Similarly, kidney sections of control animals exhibited comparable normal histological features, with no detectable abnormalities. The absence of pathological alterations in both groups suggests that *B. cepacia* infection, under the conditions tested, did not elicit detectable renal involvement or compromise kidney integrity across the study duration (**Figure 1** and **Figure 2**).

### Differential IL-18 and IL-1β responses in plasma and tissue lysates post-infection with *B. cepacia*

Next, we sought to determine the levels of IL-18 and IL-1β using ELISA were performed with both plasma and tissue lysates. The results shown that IL-18 concentration in plasma samples increased steadily from control to day 3 and reached its peak on day 7, and then declining on day 14 and 21, dropping below the baseline although not significant (p=0.08). In tissue lysates, there was a significant decline in IL-18 from control to day 3 post-infection (p<0.05). Furthermore, there was a slight increase in IL-18 which declined on day 21 (**Figure 3**). Similarly, IL-1β levels in plasma increased from control to day 3 reaching its peak on day 7, and declining on day 14 and 21, dropping below the control concentration (p=0.06). In tissue lysates, IL-1β levels witnessed remarkable decline on day 3 post-infection as compared to the control group (**Figure 3**). Hence, it was evident that IL-1β production was suppressed during the early phases of experimental *B*.*cepacia* infection.

**Figure 3.**
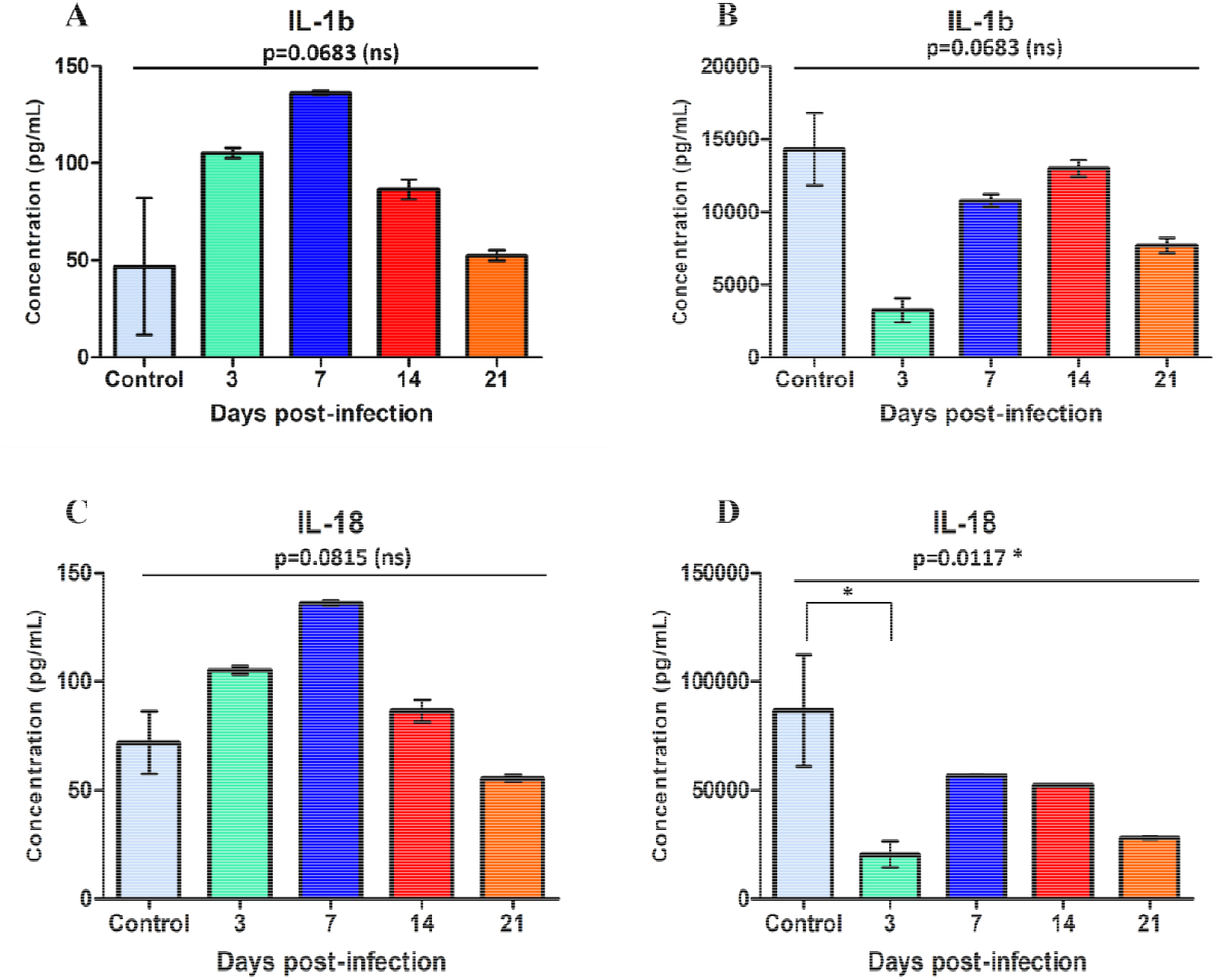
Plasma IL-1β and IL-18 concentrations across the experimental groups following infection of BALB/c mice with B. cepacia. **A)** Bar graph showing the concentration of IL-1β (pg/mL) in control and at 3, 7, 14, and 21 days post-infection. IL-18 levels increase progressively, peaking on day 7, followed by a gradual decline through days 14 and 21. **B)** The concentration of IL-18 (pg/mL) in control and day 3, 7, 14, and 21 post-infection. IL-1β levels peak early on day 3, decrease at day 7, exhibiting a partial rebound on day 14, and witnessing a decline again on day 21.

### Temporospatial alterations in plasma cytokine landscape following *B. cepacia* infection of experimental BALB/c mice

With respect to plasma cytokine levels, there was an increase in the concentrations on day 3 post-infection, whereas IL-5 levels were significantly increased as compared to other infected groups as well as control mice. In addition, almost all the cytokines increased on day 3. No significant differences were observed in IL-2 and IL-4 concentrations between the infected and control groups (p=0.6 and p=0.2). IL-2 concentrations were stable throughout the experiment, although IL-4 levels showed a slight increase on day 21 compared to the control (**Figure 3** and **Figure 4**). A transient significant increase in IL-5 was observed during day 3 post-infection as compared to the control. In contrast, IL-5 levels at later time-points were declined and were close to the baseline values (p<0.05). The concentrations of IL-10, IFN-γ and GM-CSF showed slight changes without any significance between the infected and control samples (p=0.2, p=0.4 and p=0.1, respectively). The levels of TNF-α showed a significant elevation on day 3 post-infection relative to the control group but at subsequent time-points, the levels decreased as compared to the control group (p<0.05). The concentration of IL-5 and TNF-α were significantly increased during day 3 post-infection as compared to the control mice although this occurred only for a shorter duration before falling near the baseline value.

**Figure 4.**
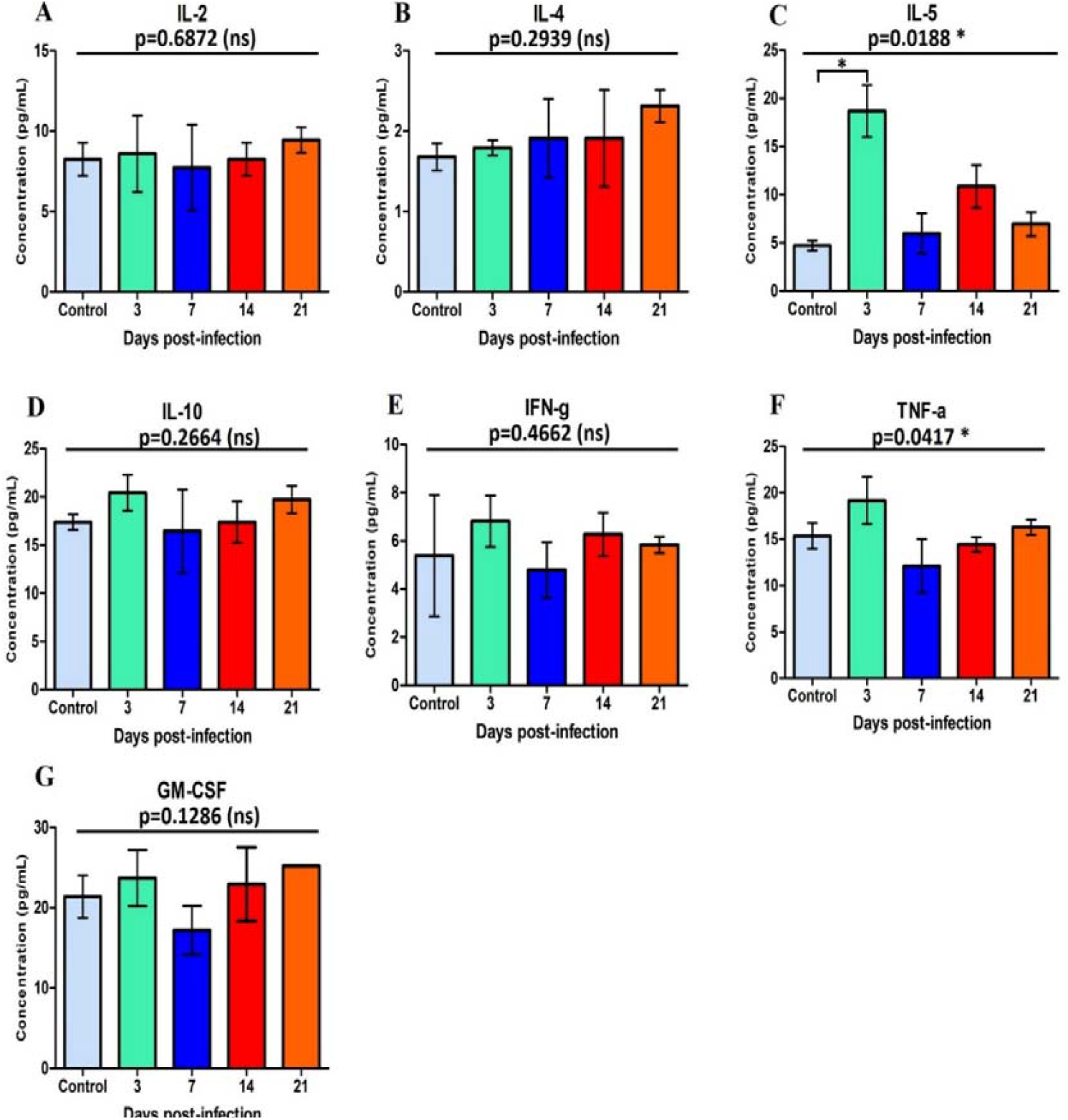
Plasma cytokines concentration across the experimental groups following infection of BALB/c mice with B. cepacia. **A** - **G)** Bar graph showing the concentration of IL-2, IL-4, IL-5, IL-10, IFN-γ, TNF-α, and GM-CSF (pg/ml) in control and on day 3, 7, 14 and 21 post-infection animals.

### TNF-α levels were significantly higher in the tissue lysates of *B*.*cepacia*-infected BALB/c mice

Multiplex cytokine analysis on the tissue lysates showed very significantly higher abundance of TNF-α in infected mice compared to control (p<0.05), whereas the levels of IL-2, IL-4, IL-5, IL-10, IFN-γ and GM-CSF were relatively lower. The levels of IL-2 in the infected group were slightly reduced as compared to the control across all time points. However, there was a significant decrease in the level of IL-2 on day 21 (p<0.05). IFN-γ levels also witnessed significant decline in infected mice (p<0.05). There was significant decrease of IFN-γ from day 14 to 21 (**Figure 4**). Cytokine concentrations of IL-4, IL-5, IL-10 and GM-CSF in infected group were either lower than or equal to the concentration observed in the control group (**Figure 5**). In summary, the cytokine profile in tissues shows that experimental *B. cepacia* infection leads to local inflammation through the TNF-α responses, although there appears to be impairment of other immune responses involving adaptive cytokines.

**Figure 5.**
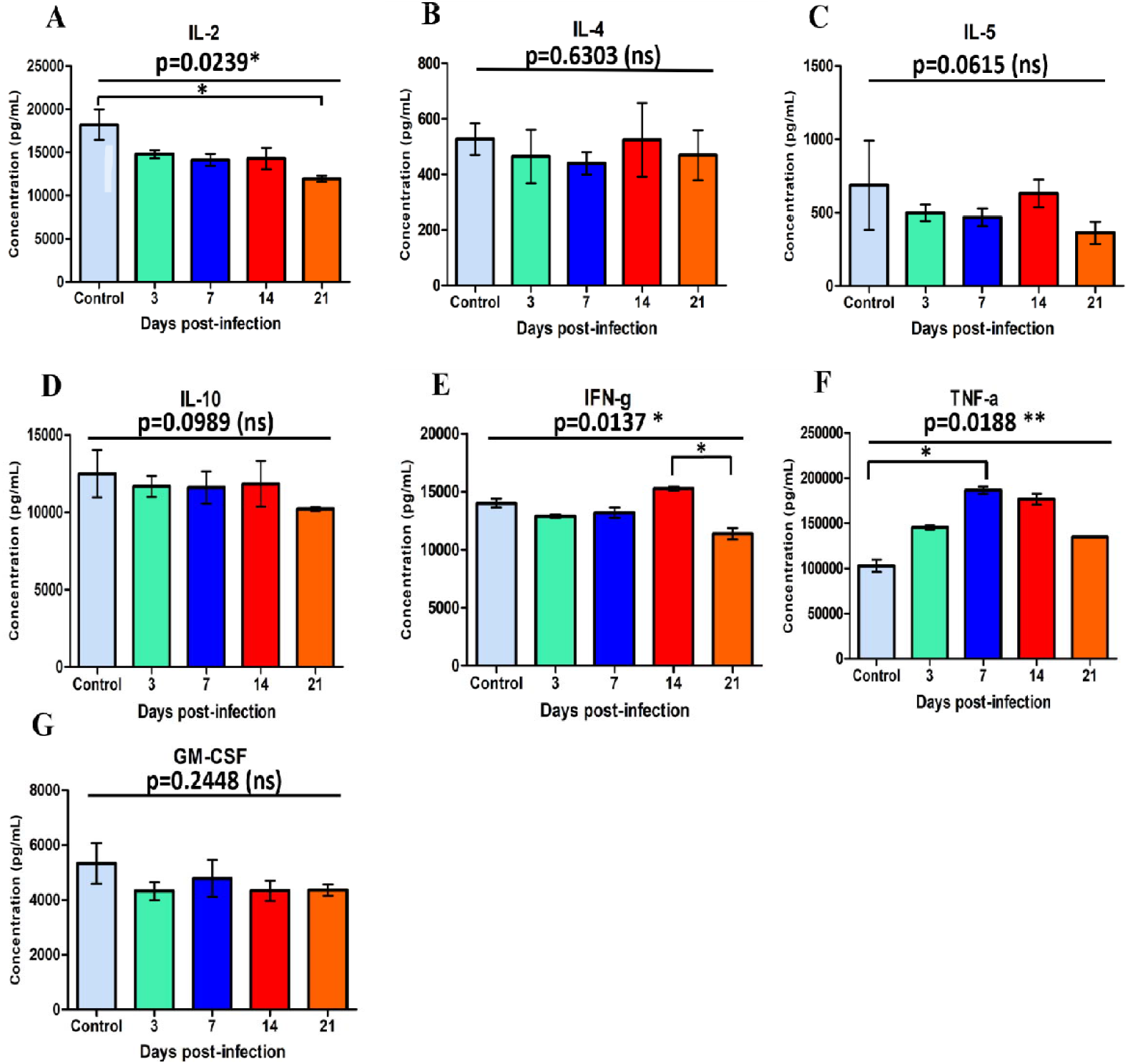
Tissue cytokines concentration across the experimental groups following infection of BALB/c mice with B. cepacia. **A** - **G)** Bar graph showing the concentration of IL-2, IL-4, IL-5, IL-10, IFN-γ, TNF-α, and GM-CSF (pg/ml) in the control and day 3, 7, 14 and 21 post-infection groups.

### Time-dependent modulation of serum MMP-2 activity was evident during *B. cepacia* infection in experimental BALB/c mice

Next, we aimed to evaluate plasma MMP-2 activity to assess infection-associated proteolytic changes during *B. cepacia* infection in the experimental BALB/c mice. Plasma of control and infected BALB/c mice at days 3, 7, 14, and 21 post-infection were analyzed by gelatin zymography under non-reducing conditions following separation on SDS-PAGE gels co-polymerized with gelatin (**Figure 6**). Our analysis demonstrated detectable MMP-2 gelatinolytic activity across all the experimental groups following infection. Importantly, variations in band intensity were observed across different time-points, suggesting dynamic regulation of MMP-2 activity over the infection course. Together, our findings indicate a time-dependent modulation of proteolytic activity associated with *B. cepacia*-induced inflammatory responses in the infected BALB/c mice.

**Figure 6.**
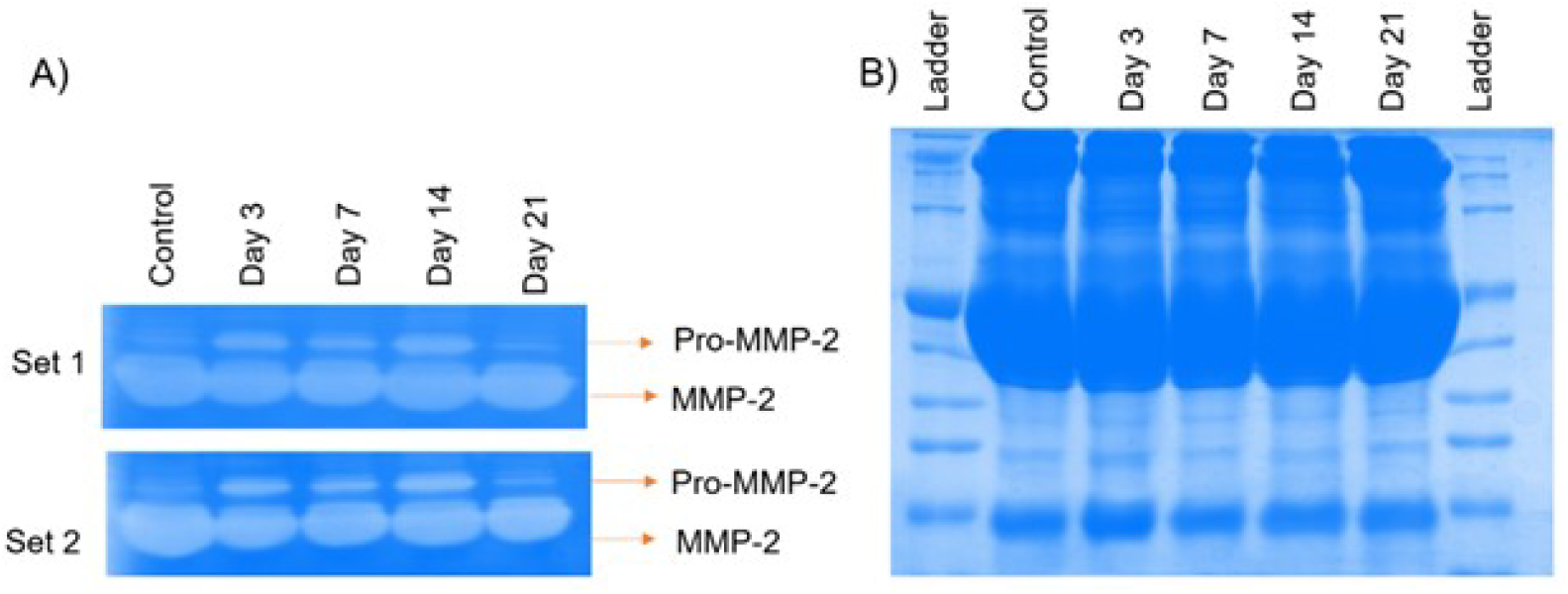
Gelatin zymography analysis of MMP-2 activity in mice plasma samples and corresponding Coomassie staining for protein loading control. **A**) Gelatin zymography was performed to evaluate MMP-2 activity in plasma samples collected from mice at different time-points (Control, Day 3, Day 7, Day 14, and Day 21). Plasma were diluted (1:10) and resolved under non-reducing conditions on gelatin-containing SDS-PAGE gels. Clear bands against the stained background indicate gelatinolytic activity corresponding to MMP-2. Two independent experimental sets (Set 1 & Set 2) are shown, demonstrating reproducibility of the observed proteolytic patterns. **B**) A parallel SDS-PAGE gel stained with Coomassie Brilliant Blue to confirm equal protein loading across all lanes. Comparable band intensities across samples indicate uniform protein input. The position of MMP-2 is indicated based on its expected molecular weight.

**Figure 7.**
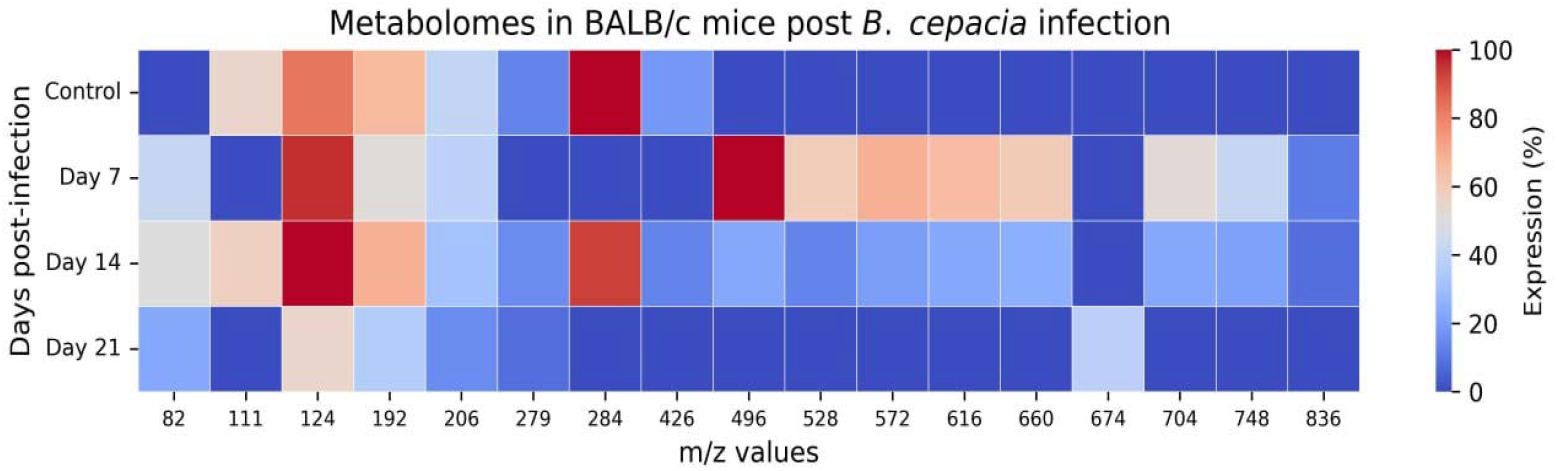
Heat map showing the relative abundance of metabolite peaks across control, day 7, day 14, and day 21 samples. During B. cepacia infection alterations in the quantity of bioactive lipid metabolites were noticeable with time. Higher levels of phospholipid and fatty acid derived metabolites were observed on day 7 and 14. Lower quantities of niacinamide, pantothenic acid, and phospotidylcholine/sphingomyline metabolites were detected on day 21 (blue marks low intensity and red high intensity).

**Figure 8.**
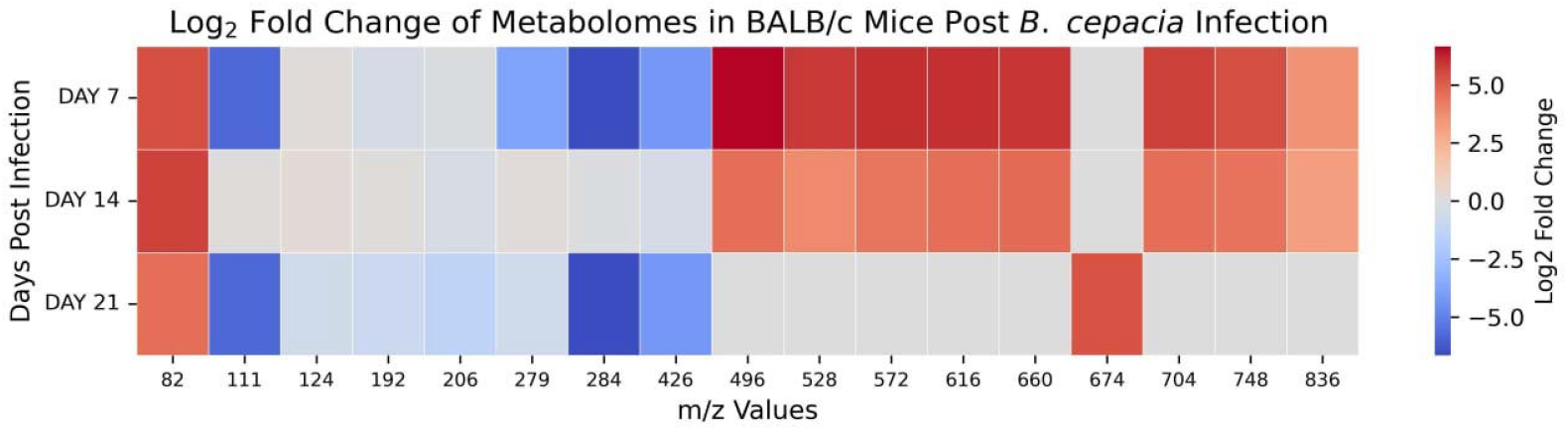
Fold change heatmap shows the differential metabolite expression in BALB/c mice following B. cepacia infection. The heat map of metabolites shows the average quantities of all metabolites from each sample (shades of blue marks low intensity and red high intensity).

**Figure 9.**
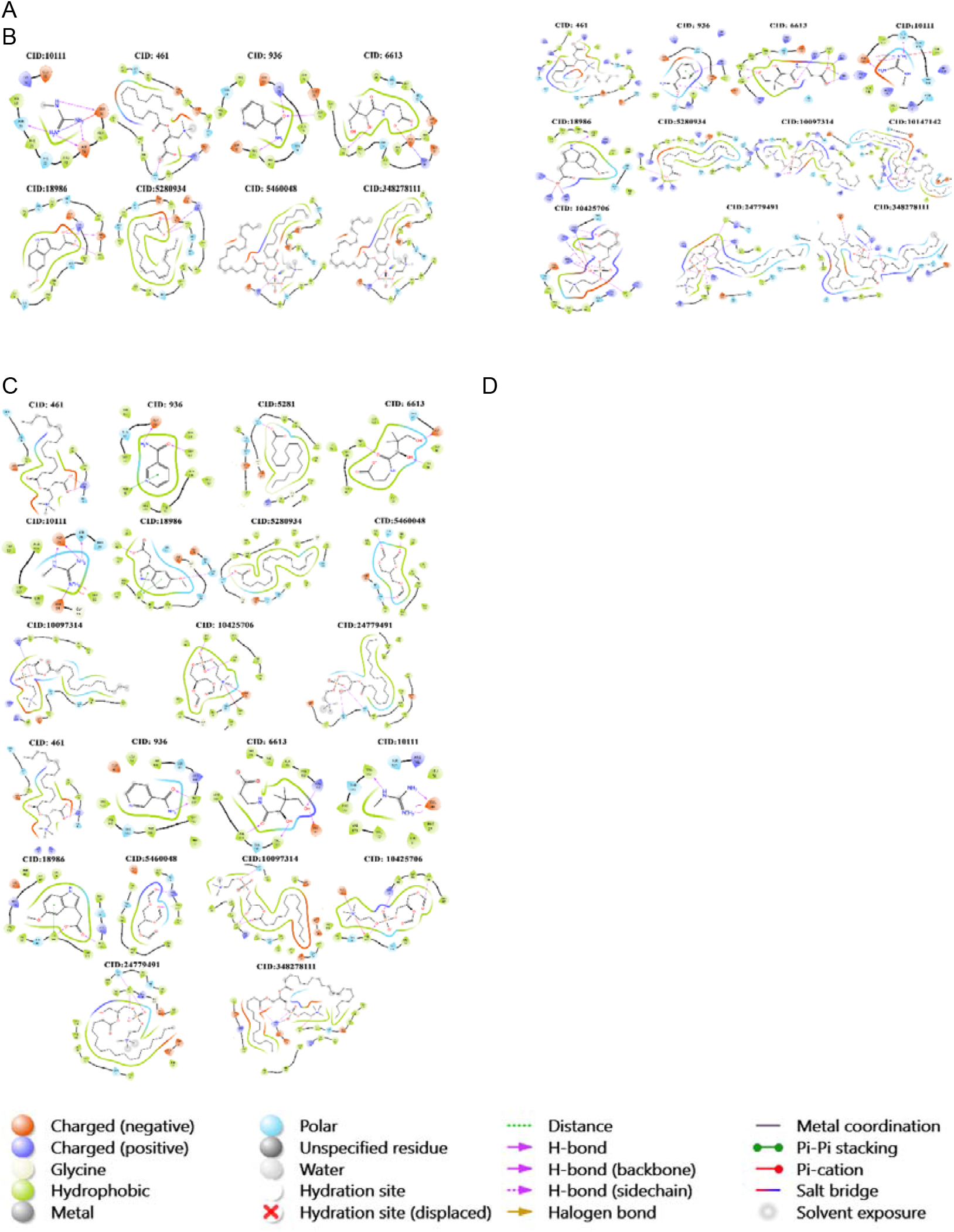
Molecular docking analysis: A) Molecular docking analysis of RNA-binding protein Hfq. 2D molecular interaction diagrams of selected compounds docked with RNA-binding protein Hfq showing hydrogen bond contacts within the active binding site. **B) Molecular docking analysis of porin-like protein BUsg_347**. 2D molecular interaction diagrams of selected compounds docked with porin-like protein BUsg_347 showing hydrogen bond contacts within the active binding site. **C) Molecular docking analysis of AHL receptor**. 2D molecular interaction diagrams of selected compounds docked with AHL receptor showing hydrogen bond contacts within the active binding site. **(D) Molecular docking analysis of AHL synthase**. 2D molecular interaction of selected compounds docked with AHL synthase showing hydrogen bond contacts within the active binding site.

### Principal component analysis shows a distinct temporal immunometobolomics profile

PCA analysis across different time points was measured for the integrated cytokine and metabolome profiles, where PC1 and PC2 account for 43.29% and 30.62% of the total variance. In the PCA analysis, we have observed distinct clustering of samples across the time period post-infection, with Day 7 showing a marked separation from other cohorts, indicative of the suggested differential immunometabolomic expression. In contrast, days 3 and 21 clustered closely, suggesting an overall cytokine and metabolomic signature, while day 14 exhibited an intermediate profile between acute and recovery stages (**Supplementary Figure 1)**

### Metabolomic landscaping of plasma revealed differential alterations following *B. cepacia* infection of BALB/c mice across different time-points

Next, we studied the plasma of *B. cepacia*-infected BALB/c mice for metabolites using untargeted high-resolution mass spectrometry (HRMS) across the different time-points. The metabolites were identified by their mass-to-charge ratio (m/z) using a positive ionization mode. Metabolomes showed differential expression across the cohorts (**Table 1**), with the majority of metabolites associated with amino acid metabolism, fatty acid metabolism, phospholipid metabolism and energy metabolism. Metabolites identified included methylguanidine, guanidine, niacinamide, pantothenic acid, 5-methoxyindoleacetate, linoleic acid derivatives, stearic acid, palmitoylcarnitine, lysophosphatidylcholine [LysoPC (16:0) and LysoPC (18:0)], sphingomyelin metabolites, metabolite species of phosphatidylcholine/sphingomyelin, phosphatidylcholine (34:0), and triacylglycerol **(Supplementary Table 3)**.

**Table 1.** Logarithmic fold change analysis of differentially expressed metabolites in BALB/c mice following *B. cepacia* infection.

| Sl. No. | Peak | Control (%) | Day 3 (%) | Day 7 (%) | Day 14 (%) | Day 21 (%) | FD7 | FD14 | FD21 | FD (14/7) | FD (21/7) | FD (21/14) |
| --- | --- | --- | --- | --- | --- | --- | --- | --- | --- | --- | --- | --- |
| 1 | 82 | - | ND | 42 | 50 | 23 | - | - | - | 0.252 | -0.869 | -1.120 |
| 2 | 111 | 55 | ND | ND | 58 | ND | - | 0.08 | - | - | - | - |
| 3 | 124 | 83 | ND | 96 | 100 | 55 | 0.210 | 0.269 | -0.594 | 0.059 | -0.804 | -0.862 |
| 4 | 192 | 66 | ND | 51 | 69 | 36 | -0.372 | 0.064 | -0.874 | 0.436 | -0.503 | -0.939 |
| 5 | 206 | 41 | ND | 39 | 32 | 15 | -0.072 | -0.358 | -1.451 | -0.285 | -1.379 | -1.093 |
| 6 | 279 | 13 | ND | ND | 15 | 8 | - | 0.206 | -0.700 | - | - | -0.907 |
| 7 | 284 | 100 | ND | ND | 93 | ND | - | -0.105 | - | - | - | - |
| 8 | 426 | 18 | ND | ND | 13 | ND | - | -0.469 | - | - | - | - |
| 9 | 496 | ND | 1 | 100 | 23 | ND | - | - | - | -2.120 | - | - |
| 10 | 528 | ND | ND | 59 | 13 | ND | - | - | - | -2.182 | - | - |
| 11 | 572 | ND | ND | 69 | 20 | ND | - | - | - | -1.787 | - | - |
| 12 | 616 | ND | ND | 65 | 23 | ND | - | - | - | -1.499 | - | - |
| 13 | 660 | ND | ND | 60 | 25 | ND | - | - | - | -1.263 | - | - |
| 14 | 674 | ND | ND | ND | ND | 38 | - | - | - | - | - | - |
| 15 | 704 | ND | ND | 52 | 23 | ND | - | - | - | -1.177 | - | - |
| 16 | 748 | ND | ND | 42 | 21 | ND | - | - | - | -1 | - | - |
| 18 | 836 | ND | 4 | 11 | 8 | ND | - | - | - | -0.459 | - | - |

Based on results from heat mapping log2 fold changes, it is evident that metabolite expressions vary between day 7, day 14 and day 21 in infected mice as compared to control. Day 7 was characterized by higher expression of lipid-related metabolites such as lysopospotidylcholine (16:0), lysophospotidylcholine (18:0), sphingomyelin fragment, phospotidylcholine (34:1), phospotidylcholine/sphingomyline, and day 21 was characterized by decrease of lysopospotidylcholine (16:0), lysophospotidylcholine (18:0), sphingomyelin fragments, phospotidylcholine (34:1) and phospotidylcholine, **(Table 1)**.

The continued decline of multiple metabolite classes at day 21 suggests that there were adaptive metabolic changes or a recovery phase during the later stages of *B*.*cepacia* infection. The appearance of other unique metabolite classes such as phospotidylcholine/sphingomyline (m/z 674) was evident on day 21. Overall, the fold change analysis showed that lipid and fatty acid metabolism and inflammatory pathways are altered in *B. cepacia* infection.

### Molecular docking analysis

Based on molecular docking analysis, compounds selected had comparable binding interactions with the target proteins, which were: RNA-binding protein Hfq, porin-like protein BUsg_347, AHL receptor, and AHL synthase. Many showed strong binding affinities and stable hydrogen bond formations within the active binding sites. Of the compounds screened, CID:18986 (5-methoxyindolacetate) **(Supplementary Table 4** and **Supplementary 5)** had the highest affinity for both the Hfq and the porin-like protein BUsg_347 and CID:24779491 (lysophosphatidylcholine (18:0) **(Supplementary Table 6)** had the highest affinity for the AHL receptor. CID:10097314 (lysophosphatidylcholine (16:0) **(Supplementary Table 7)** and CID:6613 (panthenic acid) **(Supplementary Table 7)** also had the highest affinity for AHL synthase. The amino acid residues that played a significant role in stabilising the ligands were LYS18, GLU76, ARG158, ASP137, TRP62, ASP75, PHE103 and GLU40, which were involved in protein-ligand recognition process. Most of the docked complexes had hydrogen bond distances of <2.5Å and were confirmed to produce stable binding interactions in their respective receptor cavities. Together, our results suggest that the selected compounds have good potential to inhibit their respective quorum-sensing-related target proteins.

## DISCUSSION

BALB/c mice were infected intranasally with *B cepacia*, and an inflammatory response occurred in a time-dependent manner, with progressive acute inflammatory changes. Early changes occurred in the lungs, liver, and spleen although the kidneys remained relatively unaffected. We also observed that despite severe organ pathology, there was no significant loss in body weight. The lung had the greatest severity of inflammation as determined by histopathological changes (e.g., progressive interstitial pneumonitis, bronchitis, increased thickness of the alveolar septa, dense neutrophilic and mononuclear infiltration, peri-bronchial inflammation, and gradual distortion of the alveolar architecture), which are consistent with the pulmonary tropism of *Burkholderia* sp. and the chronic respiratory inflammation observed in previous murine models (Chu et al., 2002; D. P. Speert et al., 1999b). All of these findings are also consistent with those reported by others (Burns et al., 1996) who demonstrated intracellular survival of *B. cepacia* within respiratory epithelial cells and macrophages, and with Al-Khodor et al., 2014, who showed that *B. cenocepacia* escapes into the macrophage cytosol and subverts autophagy, providing mechanistic explanations for the chronic pulmonary persistence and resistance to immune clearance observed herein. The pulmonary lesions also closely resemble human *B. cepacia* pneumonia described by others (Belchis et al., 2000), with necrotising bronchopneumonia, neutrophilic exudation and alveolar septal thickening, and mirror findings in Cftr^−^/^−^ mice (Uma Sajjan et al., 2001), which reported severe bronchopneumonia, bronchiolar inflammation, PAS-positive mucus hypersecretion and prominent neutrophilic and macrophage infiltration.

In addition to pulmonary pathology, progressive hepatic injury was evident from day 3 onwards, with focal necrosis, inflammatory infiltrates and multifocal microabscesses indicating ongoing hepatocellular damage and systemic dissemination, while the spleen showed red pulp expansion, vascular congestion and progressive neutrophilic infiltration, consistent with sustained innate immune activation and bacterial persistence in reticuloendothelial organs. These hepatic and splenic lesions are consistent with experimental *Burkholderia* infections, particularly *B. pseudomallei* models, where multifocal necrotising hepatitis, inflammatory abscesses and splenic inflammatory lesions are driven by neutrophil-mediated tissue injury (Nelson et al., 2023). Others have identified spleen as a major site of *B. cepacia* persistence in BALB/c mice (D. P. Speert et al., 1999). The predominance of neutrophils in the liver and spleen supports (Nauseef, 2007) the observation that neutrophils are the principal inflammatory cell population contributing to tissue injury in *Burkholderia* infections, and reinforces the concept that neutrophil-dominated inflammation rather than acute sepsis, characterises prolonged *B. cepacia* infection. In contrast, kidneys remained histologically intact with preserved glomerular and tubular architecture and no inflammatory infiltrates, necrosis or congestion, indicating selective organ tropism of *B. cepacia* for pulmonary and reticuloendothelial tissues suggesting that renal involvement is minimal unless severe septicemia develops.

The combined use of H&E and PAS staining improved the histopathological assessment by providing a detailed overview of inflammatory infiltrates, tissue necrosis and distortion, cell injury (via H&E), as well as identification of polysaccharide-rich tissues (via PAS), such as mucous and glycoprotein deposition related to *Burkholderia* pulmonary disease (Uma Sajjan et al., 2001). Ultimately, this study has established a reliable, temporally defined, mouse histopathological model of *B. cepacia* infection in BALB/c mice that mimics chronic pulmonary and systemic inflammatory disease with little systemic morbidity. Our results corroborate previously published data that indicate that *B. cepacia* avoids immune clearance, remains intracellular in epithelial cells and macrophages, and causes prolonged neutrophil-mediated lung, liver and spleen tissue injury (Al-Khodor et al., 2014; Chu et al., 2002; D. P. Speert et al., 1999). In addition, this model provides an excellent platform for the study of the pathogenesis of *Burkholderia*, the host immune response and the evaluation of therapeutic responses to bacteria that persist and cause chronic inflammatory damage.

Our results also show considerable changes in inflammatory and immune-regulating cytokines after *B*.*cepacia* infection. We infer that *B. cepacia* infection induces an abnormal immune response in the host, since it activates inflammatory signalling while inhibiting certain other key mediators (Amemiya et al., 2020; Koo and Gan, 2006; Uma Sajjan et al., 2001). We observed that infected samples had higher levels of TNF-α and IL-4 compared to the control, whereas IL-2, IL-5, IL-10, IFN-γ and GM-CSF were found in lower levels. In addition, there were significant differences in the expression of IL-18 and IL-1β in the infected samples (Rutter et al., 2016) (Leng et al., 2008). IL-1β is a key pro-inflammatory cytokine that are produced mainly through pyroptosis, neutrophil recruitment, and stimulation of antibacterial immune responses. Therefore, the decline in the IL-1β concentration in both plasma and tissue lysate points to the loss of inflammasome-driven immune signalling in *B. cepacia* infection. Previous studies on the species of *B. cepacia* complex show their ability to suppress the host’s immune system by altering inflammasome activation (Sousa et al., 2011). In addition, decreased IL-18 production reveals dysregulated immune responses based on inflammasome activation. The role of IL-18 is well-known for its involvement in the stimulation of the IFN-γ secretion and development of Th1-mediated cellular immunity. The observed drop in IL-18 appears to have an effect on attenuated IFN-γ secretion and macrophage activation. Meanwhile, the lowered IFN-γ concentration confirms the relationship between dysregulated IL-18 signalling and impaired Th1 immunity.

On the other hand, high TNF-α levels indicate the activation of acute inflammatory change in *B. cepacia* infection. TNF-α is secreted mainly by macrophages after bacterial component detection, including LPS recognition via TLR4 (CD284). Existing studies show the increase in TNF-α, which has been associated with inflammation-induced tissue injury as well as severity of *B. cepacia* infection in patients (Mahenthiralingam et al., 2005). Therefore, the presence of high TNF-α levels may be regarded as the activation of the inflammatory process both locally and systemically.

IL-4 in the infected group was low and closely comparable with that of the control mice. IL-4 skews Th2 polarization and the reduction in IL-4 levels indicates the absence or minimal activation of anti-inflammatory pathways during *B. cepacia* infection (Koo and Gan, 2006). The decrease in IL-2 and IFN-γ indicates the inhibition of immune responses, which are responsible for pathogenic clearance. Both IL-2 and IFN-γ play vital roles in T-cell proliferation and activation, macrophage activation, and intracellular bacterial death, respectively. The reduced expression of IL-2 and IFN-γ can negatively affect bacterial clearance contributing to persistence. Evidence suggests that *B. cepacia* can persist intracellularly due to the inhibition of immune responses and suppression of macrophage activation (Uma Sajjan et al., 2001; Schwager et al., 2013). A similar conclusion can be drawn from decreased levels of IL-10 and GM-CSF. IL-10 is an immunoregulatory cytokine that prevents tissue damage while GM-CSF promotes macrophage and neutrophil functions. Therefore, the suppression of both cytokines might contribute to acute inflammation (Moore et al., 2001).

Our findings demonstrate that infection with *B. cepacia* results in a rapid innate immune response, characterized by a transient increase in IL-18, IL-1β, TNF-α, and IL-5. In contrast, several other cytokines such as IL-2, IFN-γ, IL-10, and GM-CSF remained either uninduced or poorly induced. Therefore, these observations suggest an early innate inflammatory response; however, adaptive immunity and immune regulation remain compromised as *B. cepacia* interferes with this process (Amemiya et al., 2020; Koo and Gan, 2006). Based on the results obtained, it can be concluded that infection with *B. cepacia* is associated with dysregulation of host immune activation, which involves induction of strong innate immunity responses followed by suppression/modulation of adaptive immunity. Elevated levels of innate pro-inflammatory cytokines such as IL-1β, IL-18, and TNF-α, together with the reduced induction of IL-2, IFN-γ, IL-10, and GM-CSF, suggest that *B. cepacia* may interfere with the normal progression of protective immune mechanisms.

Gelatin zymography revealed that both pro-MMP-2 and active MMP-2 had gelatinolytic activity in BALB/c mice that were infected with *B. cepacia*, which signifies that host proteolytic responses are altered by bacterial infection (Page-McCaw et al., 2007a). Comparatively high levels of MMP-2 band intensity were seen on day 7 and 14 after infection, indicating that MMP-2 activity is dynamically regulated as the disease progresses with respect to the remodelling of ECM (Elkington and Friedland, 2006a). Increased levels of MMP-2 activity in response to bacterial infection may aid in the migration of leucocytes and infiltration of inflammatory cells into the site of infection through degradation of the ECM, thereby aiding in host immune defence mechanisms (Parks et al., 2004). Elevated levels of MMP-2 in response have been associated with tissue remodelling, recruitment of inflammatory cells, and advances in pulmonary pathology (Greenlee et al., 2007). Long-term increases in MMP-2 activity may ultimately result in the development of tissue injury and/or inflammatory pathology due to tissue degradation caused by excessive ECM breakdown (Page-McCaw et al., 2007b). The simultaneous detection of both pro-MMP-2 and active MMP-2 suggests that infection activates the degradation of the ECM via proteolytic pathways during *B. cepacia* infection (Elkington and Friedland, 2006b).

Our HRMS-based metabolomic profiling of *B. cepacia-infected* BALB/c mice revealed significant metabolic changes in the following: lipids, amino acids, phospholipid remodelling, and the host energy metabolism (Gong et al., 2023). Previous studies found similar metabolic changes associated with bacterial-mediated inflammation (Ahn et al., 2017). Substantial differences in levels of lysophosphatidylcholine (LysoPC), phosphatidylcholine (PC), sphingomyelin, fatty acid derivatives, and acylcarnitine metabolites have also occurred with infection-related changes (Law et al., 2023a). Furthermore, increases seen in lysophosphatidylcholine (16:0) and (18:0) were observed on day 7 and 14 in agreement with others (Knuplez and Marsche, 2020) showing that elevated levels of lysophosphatidylcholine metabolites correlate with inflammatory signals, oxidative damage, leukocyte activation, and phospholipid turnover while experiencing an inflammatory response. Supporting evidence from previous lipidomic studies relating to bacteria causing inflammation and immune response has shown that phosphatidylcholine and sphingomyelin metabolic dysregulation were strongly correlated to both membrane reconstruction and inflammatory lipid signalling during infection (Law et al., 2023b). In addition, significant increases in fatty acid derivatives and triacylglycerol concentrations were seen in all intermediate stages of infection, indicating that metabolic adaptation and inflammation-induced stress occur within the host (Feingold and Grunfeld, 2000).

Molecular docking evaluations of selected compounds were performed with quorum-sensing-related target proteins in *B. cepacia*, including: (a) the RNA-binding protein Hfq; (b) a porin-like protein (BUsg_347); (c) the AHL receptor; and (d) the AHL synthase (Eberl, 2006). The results of molecular docking revealed that the compound 5-methoxyindolacetate (CID: 18986) was the most potent with strong binding interactions with both Hfq and the porin-like protein (BUsg_347). Additionally, lysophosphatidylcholine (18:0) (CID: 24779491) had a much stronger affinity for the AHL receptor than other compounds. Moreover, lysophosphatidylcholine (16:0) (CID: 10097314) and pantothenic acid (CID: 6613) had strong interactions with the AHL synthase. Significant ligand-stabilising amino acid residues (LYS18; GLU76; ARG158; ASP137; TRP62; ASP75; PHE103; GLU40) were identified from the molecular docking results via hydrogen bonding and hydrophobic interactions. Most docked complexes had hydrogen bond distances <2.5 Å and thus would represent stable protein-ligand complexes, and all the compounds tested in this molecular docking study may inhibit quorum-sensing virulence in *B. cepacia* (Skariyachan et al., 2023; Slinger et al., 2019).

In conclusion, our histopathological investigations, cytokine dysregulation, metabolomics landscaping and MMP-2 studies clearly demonstrate that *B. cepacia* infection activates coordinated inflammatory and tissue-destructive responses in BALB/c mice. Severe pulmonary and hepatic lesions with neutrophilic infiltrates were associated with elevated levels of TNF-α, IL-18 and IL-1β, indicating activation of innate inflammatory pathways. Increase in the levels of inflammatory lipid metabolites, including lysophosphatidylcholine [LysoPC (16:0), LysoPC (18:0)], phosphatidylcholine, sphingomyelin fragments, palmitoylcarnitine, fatty acid derivatives, and triacylglycerol observed between days 7 and 14 of infection, indicates increased turnover of phospholipids, oxidative stress, and membrane remodelling associated with active infection. In addition, elevations in pro-MMP-2 and active MMP-2 during infection indicate that extracellular matrix degradation and inflammatory cell migration occur in conjunction with tissue remodelling. Conversely, decreased levels of IL-2, IFN-gamma, IL-10, and GM-CSF are indicative of a lack of adaptive immune responses and persistent likely survival of bacteria. A gradual decline in the number of inflammatory metabolites and MMP-2 activity on day 21 of infection suggests partial metabolic adaptation and repair following tissue injury attributable to infection. Hence, our findings highlight organ-specific pathologic progression and sustained inflammation providing key insights into host–pathogen interactions.

## Acknowledgements

The authors would like to acknowledge Ms. Narmada, Department of Biotechnology, Central University of Tamil Nadu, Thiruvarur, for her valuable suggestions during the standardization of animal experiments. Prof. Vikas Gautam, Department of Microbiology, PG Institute of Medical Education and Research, Chandigarh, India, is gratefully acknowledged for providing *Burkholderia cepacia* as a generous gift. The authors acknowledge technical support extended by the High-Resolution Mass Spectrometer Facility, Vellore Institute of Technology, Vellore, India. The authors also like to acknowledge Dr. S. Rajalakshmi (Karpagam Academy of Higher Education, Coimbatore, India) for her support with the gelatin zymography assay. The authors are also grateful for the Animal House Facility, Annamalai University, Chidambaram, India. The authors also thank the Department of Science and Technology (DST), Government of India, for the financial support extended under the DST-FIST Program (Grant No. SR/FST/LS-1/2020/641) to the Department of Biotechnology, Central University of Tamil Nadu, Thiruvarur, India.

## Funding

S. R. is funded by a GAT-B Fellowship, Department of Biotechnology; A.R.A is funded by NFOBC-NBCFDC of the Ministry of Social Justice, Government of India, Senior Research Fellowship (Ref. No. 221610155057); M. A. is supported by a University Intramural Fellowship, Central University of Tamil Nadu, Thiruvarur, India.

## Competing interest

The authors have no competing interests.

## Notes

### Competing Interest Statement

The authors have declared no competing interest.

### Summary of Updates

First names and last names of authors correctly updated.Figures previously not cited in the text added now and supplementary files that were missing have now been incorporated.

