## Supplementary Table 1 for "Intranasal administration of *Burkholderia cepacia* promotes progressive acute inflammatory changes in experimental BALB/c mice"

**Supplementary Table 1:** Phenotypic and biochemical characteristics of *B. cepacia* 20209

| **Phenotypic/Biochemical test** | **Observation** | **Result** |
| --- | --- | --- |
| Grams staining | Pink/Red coloured rod | Gram negative rods |
| Nutrient agar medium | Growth | Non-fastidious |
| MacConkey agar medium | Yellow colonies | Non-lactose fermenting |
| Cetrimide agar medium | No growth | Inability to grow |
| Tryptic soy agar medium | Growth observed | Non-fastidiousness |
| Catalase test | Bubble formation | Positive |
| Blood agar | No clear zone of hemolysis | Non-hemolytic |
| Bile esculin test | No colour change | Negative |
| IMViC | (Negative, Negative, Negative, Positive) | Indole, VP, MR negative  Citrate positive |
| Arginine dihydrolase test | Color change to purple | Positive |
| Nitrate reduction test | Bubble formation, color change after addition of reagent A&B, zinc dust | Positive |
