## Supplementary Table 2 for "Intranasal administration of *Burkholderia cepacia* promotes progressive acute inflammatory changes in experimental BALB/c mice"

**Supplementary Table 2:** Characteristics of BALB/c mice cohorts

| **Mice** | | | **n=32** |
| --- | --- | --- | --- |
| **S.No** | **Cohort** | **Total no. of mice** | **Weight** |
| 1 | Control | 8 | 28 ± 2.23 |
| 2 | Day 3 | 6 | 30 ± 3.03 |
| 3 | Day 7 | 6 | 28 ± 3.5 |
| 4 | Day 14 | 6 | 29 ± 2.1 |
| 5 | Day 21 | 6 | 27.5 ± 3.4 |
