## Supplementary Table 3 for "Intranasal administration of *Burkholderia cepacia* promotes progressive acute inflammatory changes in experimental BALB/c mice"

**Supplementary Table 3:** Metabolites identified in the plasma samples

| **S. No.** | **Peak** | **Tentative metabolite** |
| --- | --- | --- |
| 1 | 82.0401 | Methylguanidine |
| 2 | 111.0075 | Guanidine |
| 3 | 124.0738 | Niacinamide |
| 4 | 192.126 | Panthenic acid |
| 5 | 206.0848 | 5-Methoxyindolacetate |
| 6 | 279.0848 | Linoleic acid derivatives |
| 7 | 284.3224 | Stearic acid |
| 8 | 426.338 | Palmetoylearnnitine |
| 9 | 496.3381 | Lysophospotidylcholine (16:0) |
| 10 | 528.4132 | Lysophospotidylcholine (18:0) |
| 11 | 572.4409 | Sphingomylin fragment |
| 12 | 616.4699 | Phospotidylcholine/lysophospholipids |
| 13 | 660.4984 | Phospotidylcholine (34:1) |
| 14 | 674.5101 | Phospotidylcholine/Sphingomyline |
| 15 | 704.5271 | Phospotidylcholine/Sphingomyline |
| 16 | 748.5562 | Phospotidylcholine (34:0) |
| 17 | 836.2759 | Triacylglycerol |
