## Supplementary Table 4 for "Intranasal administration of *Burkholderia cepacia* promotes progressive acute inflammatory changes in experimental BALB/c mice"

Molecular docking interactions of selected compounds with the Hfq protein regarding their amino acids' interactions, hydrogen bond lengths, and Glide scores.

| **S. No** | **PubChem ID** | **Ligands** | **Amino Acid** | **Bond Length (Å)** | **Glide Score (kcal/mol)** |
| --- | --- | --- | --- | --- | --- |
| 1 | CID:10111 | Methylguanidine | ASP 74  ASP 74  GLU 76  GLU 76  ASN 70 | 1.61  2.12  1.61  2.11  1.80 | -2.181 |
| 2 | CID: 461 | Palmetoylearnnitine | ASN 14  LYS 18 | 1.93  1.89 | -2.318 |
| 3 | CID: 936 | Niacinamide | PRO 11  LYS 18  ALA 77 | 2.04  1.64  2.32 | -3.388 |
| 4 | CID: 6613 | Panthenic acid | PRO 11  LYS 18  GLU 76 | 2.02  1.79  1.79 | -3.888 |
| 5 | CID:18986 | 5-methoxyindolacetate | LYS 18,  GLU 76,  ALA 77 | 1.59  2.11  1.96 | -4.360 |
| 6 | CID:5280934 | Linolenic acid derivatives (alpha) | LYS 18  ALA 77  GLU 76 | 1.70  2.37  1.86 | -3.368 |
| 7 | CID: 5460048 | Triacylglecerol | LYS 18  GLU 76 | 1.85  2.27 | -1.857 |
| 8 | CID:348278111 | Phosphatidylcholine (34:1) | LYS 18  ASN 14 | 2.13  2.22 | -1.726 |
