## Supplementary Table 5 for "Intranasal administration of *Burkholderia cepacia* promotes progressive acute inflammatory changes in experimental BALB/c mice"

Molecular docking interactions of selected compounds with the porin-like proteins on their amino acid interactions, hydrogen bond lengths, and glide scores.

| **S.No** | **PubChem ID** | **Ligands** | **Amino Acid** | **Bond Length (Å)** | **Glide Score (kcal/mol)** |
| --- | --- | --- | --- | --- | --- |
| 1 | CID:461 | Palmetoylearnnitine | ARG 158  ARG 158  ARG 106  ARG 106  MET 1 | 2.08  1.66  2.27  2.22  1.95 | -3.24828 |
| 2 | CID:936 | Niacinamide | ARG 106  ASN 3  HIS 130 | 2.11  2.13  2.08 | -3.51339 |
| 3 | CID:6613 | Panthenic acid | ARG 158  ARG 106  ARG 106  MET 1  LYS 2  VAL 139  ILE 142  TYR 141 | 2.03  2.05  2.12  1.93  2.63  1.79  2.39  2.24 | -4.56529 |
| 4 | CID:10111 | Methylguanidine | ASP 137  ASP 137  SER 134 | 1.94  2.16  2.19 | -3.34821 |
| 5 | CID:18986 | 5-methoxyindolacetate | VAL 139 | 2.12 | -7.54697 |
| 6 | CID:5280934 | Linolenic acid derivatives (alpha) | ARG 106  ARG 106  MET 1 | 2.71  1.68  2.24 | -4.76141 |
| 7 | CID:10097314 | Lysophosphatidylcholine (16:0) | LYS 87  ARG 158  ARG 106 | 1.97  2.13  2.66 | -2.87939 |
| 8 | CID:10147142 | Phosphatidylcholine (34:0) | LYS 87  LYS 87  ARG 106  ARG 158  ARG 158  ASN 151 | 2.26  1.90  1.81  2.19  2.19  2.54 | -3.23303 |
| 9 | CID:10425706 | Phosphatidylcholine/Lysophospholipids (+1) | MET 1  ARG 106  ARG 106  ARG 106  ARG 158  ARG 158  LYS 87 | 2.16  1.96  1.60  2.17  1.82  1.79  1.85 | -4.54854 |
| 10 | CID:24779491 | Lysophosphatidylcholine (18:0) | MET 1  ALA 149  ARG 158  ARG 158  LYS 87  ARG 106  ARG 106  ARG 106 | 2.20  2.42  2.16  2.30  1.78  2.32  2.18  2.06 | -2.60956 |
| 11 | CID:348278111 | Phosphatidylcholine (34:1) | LYS 71  LYS 113 | 1.74  2.27 | -1.78213 |
