## Supplementary Table 6 for "Intranasal administration of *Burkholderia cepacia* promotes progressive acute inflammatory changes in experimental BALB/c mice"

Molecular docking interactions of selected compounds with the AHL on their amino acid interactions, hydrogen bond lengths, and glide scores.

| **S. No** | **PubChem ID** | **Ligands** | **Amino Acid** | **Bond Length (Å)** | **Glide Score (kcal/mol)** |
| --- | --- | --- | --- | --- | --- |
| 1 | CID:461 | Palmetoylearnnitine | LYS 49 | 1.71 | -2.35722 |
| 2 | CID:936 | Niacinamide | TRP 62  ASP 75 | 2.10  1.88 | -5.29626 |
| 3 | CID:5281 | Stearic acid | SER 98 | 2.04 | -5.87164 |
| 4 | CID:6613 | Panthenic acid | ASP 75  TRP 62  TYR 58 | 2.03  1.71  2.59 | -3.20324 |
| 5 | CID:10111 | Methylguanidine | SER 76  ASP 75  ASP 75 | 1.92  1.62  1.99 | -4.14799 |
| 6 | CID:18986 | 5-methoxyindolacetate | TRP 62  THR 77 | 1.98  1.92 | -5.5203 |
| 7 | CID:5280934 | Linolenic acid derivatives (alpha) | SER 98 | 1.85 | -4.75927 |
| 8 | CID:5460048 | Triacylglecerol | THR 77  TYR 58  TRP 62 | 2.43  1.86  1.95 | -3.22841 |
| 9 | CID:10097314 | Lysophosphatidylcholine (16:0) | LYS 49 | 1.86 | -2.3496 |
| 10 | CID:10425706 | Phosphatidylcholine/  Lysophospholipids (+1) | TRP 62  TYR 58 | 1.85  2.36 | -0.222961 |
| 11 | CID:24779491 | Lysophosphatidylcholine (18:0) | GLN 67  ASN 70 | 2.10  1.98 | -6.07122 |
