## Supplementary Table 7 for "Intranasal administration of *Burkholderia cepacia* promotes progressive acute inflammatory changes in experimental BALB/c mice"

Molecular docking interactions of selected compounds with the acyl-homoserine-lactone synthase regarding their amino acid interactions, hydrogen bond lengths, and glide scores.

| **S. No** | **PubChem ID** | **Ligands** | **Amino Acid** | **Bond Length (Å)** | **Glide Score (kcal/mol)** |
| --- | --- | --- | --- | --- | --- |
| 1 | CID:461 | Palmetoylearnnitine | SER 146 | 1.73 | -2.87762 |
| 2 | CID:936 | Niacinamide | PHE 103  PHE 103 | 1.96  1.94 | -3.9258 |
| 3 | CID:6613 | Panthenic acid | PHE 27  GLU 40  PHE 144  VAL 142 | 2.17  1.83  2.12  2.09 | -5.27439 |
| 4 | CID:10111 | Methylguanidine | VAL 142  GLU 40  GLU 40 | 2.04  1.25  1.25 | -2.99854 |
| 5 | CID:18986 | 5-methoxyindolacetate | PHE 103  PHE 144 | 2.34  2.29 | -3.30856 |
| 6 | CID:5460048 | Triacylglecerol | ARG 102  PHE 103 | 2.16  2.13 | -3.90028 |
| 7 | CID:10097314 | Lysophosphatidylcholine (16:0) | GLN 30  PHE 103  PHE 103 | 2.24  1.84  2.09 | -5.30727 |
| 8 | CID:10425706 | Phosphatidylcholine/  Lysophospholipids (+1) | PHE 144  PHE 103  ALA 105 | 2.22  2.55  2.13 | -1.26322 |
| 9 | CID:24779491 | Lysophosphatidylcholine (18:0) | MET 171  SER 146 | 2.04  2.00 | -1.83214 |
| 10 | CID:348278111 | Phosphatidylcholine (34:1) | LYS 41  LYS 41 | 1.69  2.25 | -1.46722 |
