## Supplementary figures and images for "Intranasal administration of *Burkholderia cepacia* promotes progressive acute inflammatory changes in experimental BALB/c mice"

### Supplementary Figure 1

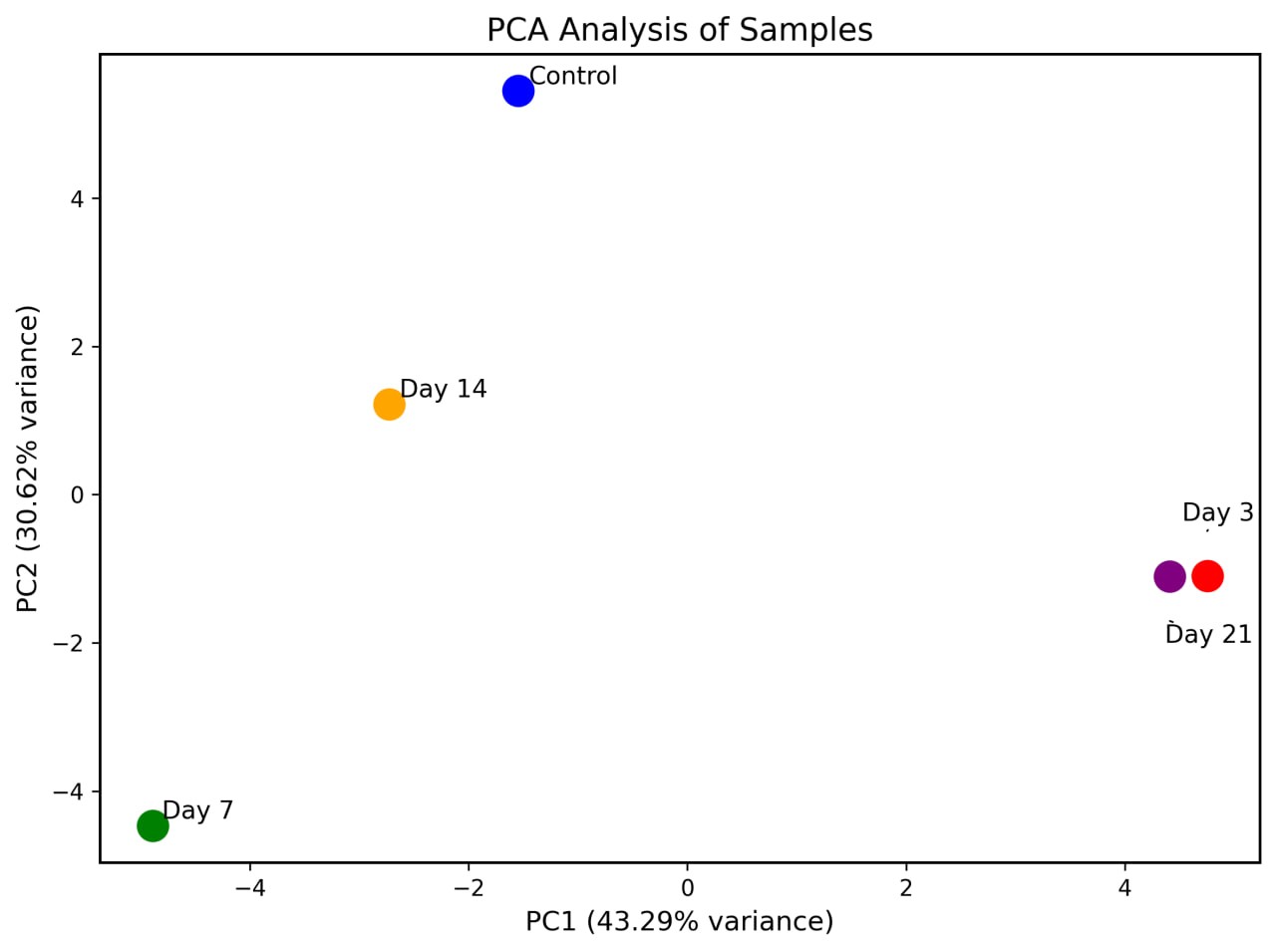
